# A ligated intestinal loop mouse model protocol to study the interactions of *Clostridioides difficile* spores with the intestinal mucosa during aging

**DOI:** 10.1101/2021.06.21.449186

**Authors:** Pablo Castro-Córdova, María José Mendoza-Leon, Daniel Paredes-Sabja

**Affiliations:** Microbiota-Host Interactions and Clostridia Research Group, Departamento de Ciencias Biológicas, Facultad de Ciencias de la Vida, Universidad Andrés Bello, Santiago, Chile; ANID-Millennium Science Initiative Program - Millennium Nucleus in the Biology of the Intestinal Microbiota, Santiago, Chile; Department of Biology, Texas A&M University, College Station, Texas, USA

**Keywords:** C. *difficile* spores, spore adherence, spore entry, aging, intestinal ligated loop, ileal loop, colonic loop, intestinal mucosa

## Abstract

The interaction of the *Clostridioides difficile* spores with the intestinal mucosa contribute to the persistence and recurrence of the infection. Advanced age is one of the main risk factors to manifest *C. difficile* infection and recurrence. However, the interaction of *C. difficile* spores with the intestinal mucosa during aging has not been evaluated. In the present work, we provide a detailed protocol with all the critical information to perform an intestinal ligated loop. Using this technique in a mouse model, we evaluated *C. difficile* spore adherence and internalization to the ileum and colonic mucosa during aging. Consequently, our data suggest that spore internalization in the ileum and colonic mucosa is higher in elderly than in adults or young mice. Also, our data suggest that spore-adherence to the ileum and colonic mucosa decreases with aging.

## INTRODUCTION

*Clostridioides difficile* is a Gram-positive, anaerobic, and spore former bacterium that is the leading pathogen causing hospital acquiring diarrhea associated with antibiotics [1, 2]. The *C. difficile* infection (CDI) is characterized by the manifestation of diarrhea that can produce mild to watery diarrhea, abdominal pain, and tenderness [3]. In severe cases, patients can become dehydrated or produce toxic megacolon [3]. CDI is lethal in ∼5% of the infected patients [4, 5]. The ∼15–30% of recovered of *C. difficile* diarrhea manifest a recurrent CDI (R-CDI) [5, 6].

The main two risk factors for CDI are the continuous alteration of the microbiota caused by antibiotics, and age over 65 years old [7, 8], being the 91% of the CDI deaths in this age group [9]. This increasing association in CDI risk with aging may be explained by age-related physiologic changes such as the immunosenescence and age-related dysbiosis of the intestinal microbiota causing a reduced the protective role against *C. difficile*. The immunosenescence is characterized by a progressive decrease in the effectiveness of the immune system associated with aging, increasing the susceptibility to infections in older adults due to the impaired innate and adaptive immune response [10]. For example, during aging occurs a dysfunctional antigen-presenting cell, reduced chemotaxis to inflammatory stimuli of natural killers, neutrophils [11, 12], reduced activity in bacterial phagocytosis by monocytes and macrophages [12, 13]. There is also a reduced antibody response to exogenous antigens and vaccines by B-cells [14]. These changes may be explained by altered intracellular communication, telomere attrition, epigenetic alterations in the earliest hematopoietic stem cells [14]. Therefore, elderly patients have an increased risk of bacterial infections such as CDI.

Age-related dysbiosis is characterized by a reduced species diversity enriched in pro-inflammatory commensals bacteria [15]. In particular by a decline of *Bifidobacterium* [16] and Clostridiales and with enrichment in Proteobacteria and an overrepresentation of Enterobacteriaceae [14]. That age-related dysbiosis is associated with a reduced protective role of the microbiota against CDI [17, 18]. For example, it has been reported that fecal emulsions from geriatrics patients have low inhibitory activity in the growth of *C. difficile in vitro* compared with fecal emulsions from healthy adults [17].

During CDI, *C. difficile* forms metabolically dormant spores that are essential for R-CDI [19]. Accordingly, with this observation, we developed a surgical procedure of intestinal ligated loop, which allowed us to demonstrate that *C. difficile* spores are able to adhere and internalize into the intestinal mucosa contributing to disease recurrence [20]. However, whether aging affects the adherence and internalization of *C. difficile* spores to the intestinal mucosa remains unclear. Due to the lack of protocols describing the intestinal ligated loop in detail, we decided to provide this step-by-step protocol that brings information on critical points to perform ligated ileum and colonic loop injected with *C. difficile* spores. Also, this work provides information on the processing and mounting of the tissues to acquire high-resolution confocal images and quantify the spore adherence and internalization to the intestinal mucosa. Next, using this technique, we evaluate the spore adherence and internalization into the intestinal mucosa of young (7-weeks-old), adult (1-year-old), and elderly mice (2-years-old). Our results suggest that spore adherence decrease with the aging in both ileum and colonic mucosa. However, suggest that the spore entry is increased in elderly mice to the intestinal mucosa.

## 2.- MATERIAL

### 2.1 Reagents

Chemicals

1. Ethanol 95% (Winkler, Chile).
2. Ophthalmic ointment (Pharmatech, Chile).
3. Isoflurane USP (Baxter Healthcare, Puerto Rico).
4. 10% Povidone-iodine solution (DifemPharma, Chile).
5. Triton 100-X (Merck, Germany).
6. Sucrose (Winkler, Chile).
7. Paraformaldehyde (Merck, Germany).
8. Dako fluorescent mounting medium (Dako, USA).
9. Bovine derum albumin (Sigma-Aldrich, USA).
10. Sodium Chloride (Merck, Germany).

Others

1. Heating pad (Imetec, Italy).
2. Extra thick blot paper (BioRad, USA).
3. Masking tape (3M, USA).
4. Disposable Razor Schick Xtreme 3 sensitive skin (Schick, USA).
5. Immersion oil for fluorescence microscopy, type LDF, formula code 387 (Cargille, USA).
6. Microscopy glass slide (Hart, Germany).
7. Cover slide (Hart, Germany).
8. Plastic Petri dish (Bell, Chile).
9. Scotch transparent tape (3M, USA).
10. Microcentrifuge 1.5mL tubes.
11. Microcentrifuge 0.5mL tubes.
12. 500mL autoclavable Glass Bottle (Schott Duran, USA).
13. 29G insulin syringe (Nipro, USA).
14. Towel paper.
15. Surgical silk suture 3/0 HR20 (Tagum).
16. Vinylic tape (Scotch 3M, USA).
17. 0.5% bleach.
18. Virkon-S (DuPont).

Material and Equipment.

1. Stainless-steel surgical tray.
2. Scalpel.
3. Watchmaker forceps.
4. Anatomical forceps, stainless steel.
5. Anatomical forceps fine, stainless steel.
6. Forceps Dumont N°5, super fine tips.
7. Dressing Forceps.
8. Surgical scissor blunt & sharp tip.
9. Iris scissors.
10. Infrared heat lamp light bulb.
11. Isoflurane chamber with a facemask for mice.
12. Orbital shaker.
13. Epifluorescent Microscope Olympus BX53.
14. Confocal microscope Leica SP8.
15. Biosafety cabinet.

### 2.2 Solutions

#### 10× Phosphate buffered saline (PBS)

Stock solution; Calculate the reagents required to prepare 1L of 1.37 M NaCl, 27 mM KCl, 100 mM Na_2_HPO_4_, 18 mM KH_2_PO_4,_ and dissolve it in 800mL of Milli-Q water. Then the pH was adjusted to 4.5 with HCl and add Milli-Q water to 1L. Sterilize by autoclaving for 20 min at 121 °C with 15 psi, and store at room temperature.

#### 1× (PBS)

Dilute 10× PBS stock solution 1:9 (100 mL of 10× PBS into 900 mL of Milli-Q water) and then sterilize by autoclaving for 20 min at 121 °C with 15 psi. Store at room temperature.

#### Saline solution

0.9% NaCl. Dissolve 9 g of NaCl in 1L of Milli-Q water. Pass through 0.45-µm filters and sterilize by autoclaving for 20 min at 121 °C with 15 psi. Store at room temperature.

#### Fixing solution

30% sucrose in PBS–4% paraformaldehyde. In the first place, a solution of PBS–4% paraformaldehyde was prepared as follows: for 1L, add 40 g of paraformaldehyde powder to 800 mL of 1× PBS. Heat to 60 °C in a fume hood (no dot boil). If it does not dissolve, raise the pH adding 5N NaOH drop by drop until a clear solution is formed. Cool the solution, adjust the pH to 8.0, and adjust the volume to 1L with 1× PBS. Pass thought 0.45-µm filter to remove particles. Aliquot in small volume and store at 4 °C for use in 1–2 weeks or store at 20 °C for up to 1 year. To prepare 100 mL of 30% sucrose in PBS–4% paraformaldehyde, 30 g of sucrose were dissolved in PBS–4% paraformaldehyde with a final volume of 100 mL.

#### Permeabilizing solution

a PBS–0.2% Triton X-100 solution was prepared as follows: a stock solution of 10% Triton X-100 was prepared in PBS. To prepare 10 mL of the stock solution, dilute 1 mL of Triton X-100 in 9mL of PBS with gentle shaking. PBS–0.2% Triton X-100 solution was made by dilution 1:49 of the stock with sterilized 1× PBS. The stock solution was stored at 4 °C.

#### Blocking solution

PBS–3% BSA solution. To prepare 10 mL of the solution, 0.3g of BSA was dissolved in 8 mL of 1× PBS, then add PBS to 10mL and sterilize by filtering with 0.2-µm syringe filter. The solution was stored at 4 °C.

## 3.- METHODS

### Mice used

7-weeks (*n* = 5), 1-year-old (*n* = 3) and 2-years-old (*n* = 4) C57BL/6 (male or female) were obtained from the breeding colony at the Departamento de Ciencias Biológicas of the Universidad Andrés Bello derived from Jackson Laboratories. Mice were housed with *ad libitum* access to food Prolab RMH 3000 (Prolab, USA) and autoclaved distilled water. Bedding and cages were autoclaved before use. Mice were housed with 12-h cycle of light and darkness, at 20–24 °C with 40–60% of humidity. All procedures complied with all ethical regulations for animal testing and research. This study received ethical approval from the Institutional Animal Care and Use Committee of the Universidad Andrés Bello.

### Biosafety and personal protection elements

During the surgery, the use of personal protective equipment is required, such as a disposable laboratory coat, goggles, gloves, cap, and mask when you are manipulating the fixing solution. The surgery is not performed under the biological safety cabinet. However, all processes that produce aerosol, such as preparing the syringe with *C. difficile* spore inoculum, opening and cleaning the tissues infected with *C. difficile* spores, were performed under the biosafety cabinet. Finally, all the used surfaces were disinfected with a solution of 0.5% bleach and 1:200 of Virkon-S.

### Surgery

All surgical procedures were performed under clean but non-sterile conditions. The surgical scissors and forceps were autoclaved before usage except the forceps Dumont N° 5 that are not autoclavable, so they were washed with soap, 0.5% bleach, and 70% ethanol.

#### Day 1

All the required materials such as isoflurane, povidone-iodine, 70% ethanol, ophthalmic ointment, heat-pad, a stainless-steel surgical tray attached to an isoflurane mask, syringe with saline solution, syringes with *C. difficile* spores, silk braided silicon suture, autoclaved towel paper, masking tape, and the surgical material as forceps, scissors, were organized in a manner that they are easily accessible to the hand (Fig 1).

**Fig 1.**
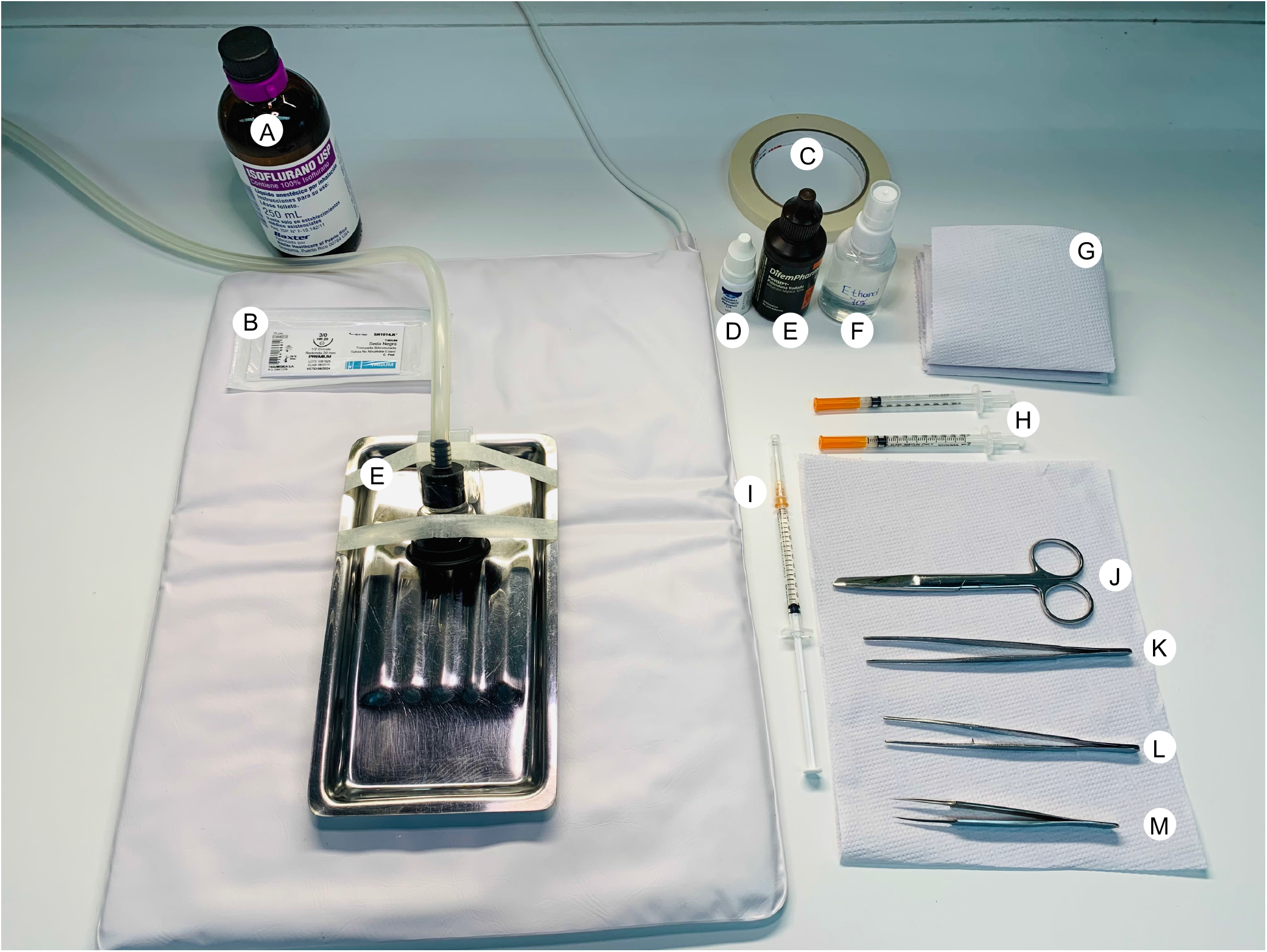
Surgical preparation for intestinal ligated loop procedure. In a clean, disinfected, but non-sterile condition, the surgical instruments were prepared as follow: (A) isoflurane-USP, (B) silk braided silicon-coated suture 3-0 HR20, (C) masking tape, (D) ophthalmic ointment, (E) 10% povidone-iodine solution, (F) 70% ethanol solution, (G) sterilized paper towels, (H) 29G insulin syringe with 100µL of 0.9% NaCl containing 5 × 10^8^ *C. difficile* spores, (I) 29G insulin syringe containing sterile 0.9% NaCl to hydrate the tissues), (J) surgical scissors Sharp/blunt, stainless steel, (K) anatomical forceps, stainless steel, (L) anatomical forceps fine, stainless steel, (M) forceps Dumont N°5, super fine tips.

1. Male or female, 18–25g mice C57BL/6 of 8–12 weeks were fasted overnight (15 h) before the surgery with free access to water.
2. Depth anesthesia was induced in ∼3min with 4% (vol/vol) isoflurane and a flow of 0.6L/min of with an isoflurane induction chamber.
3. Take out the mouse from the isoflurane induction chamber, put the mouse prone into the surgery tray, and put the snout (nose and mouth) into the isoflurane mask. The surgery bed is over a heating pad to avoid hypothermia during the procedure. Reduce the isoflurane concentration to 2% (vol/vol).
4. Add ophthalmic drops in the eyes to avoid corneal drying.
5. Turn the mouse to supine position.
6. Check anesthetic depth by non-response to hind limb toe pinch.
7. Using small pieces of masking tape, fix the limbs of the mice to the surgery bed.
8. Dampen the hair of the abdominal area with 70% ethanol either by spraying or by dipping. Clean with a paper towel. Let it dry.
9. With a disposable razor, shave the abdominal zone.
10. Clean the shaved abdominal zone with povidone-iodine and clean it with towel paper (Fig 2A). For steps 1–10, see S1 Video.
11. Using forceps and a surgical scissor sharp/blunt, perform an incision of ∼2cm in the midline of the abdominal skin (Fig 2B).
12. The skin is separated from the peritoneum using anatomical forceps.
13. Identify the linea alba, a semitransparent longitudinally white line in the peritoneum (Fig 2C).
14. Open the abdominal cavity, incise the abdominal musculature on the linea alba. For this, gently grab the musculature with anatomical forceps and retracting up. The abdominal organs are not adjacent to the muscles. Using a scissor or a scalpel makes a small opening into the peritoneal cavity in the linea alba. Extend the incision in the midline until it reaches the size of the skin cut. For steps 11–14, see S2 Video.
15. Using forceps, gently move the intestines to identify the cecum, a large J-shaped blind sac curved. Extract the cecum through the incision and identify the ileum and colon (Fig 3A). The ileum is the distal last part of the small intestine that is attached to the cavity by mesentery tissue that derives blood supply from the mesenteric artery [21]. The proximal colon is the first part of the colon that begins in the cecum.
16. When the ileum or colon has fecal material, it can be removed, pressing gently with a blunt tip of the forceps against your fingers and move it in the direction of the flow of the fecal material or to the cecum.
17. In the ileum, identify regions to be ligated where blood vessels are finely separated from the ileum’s external wall (Fig 3B). Once identified, pass the fine tip of the forceps Dumont N° 5 between the outer wall of the ileum and the blood vessels having care of not damage or puncture the blood vessels. With the tip of the forceps Domont N° 5, on the other side of the hole formed between the external wall of the ileum and the blood vessels, grasp the thread of the surgical suture, and gently pass to the other side of the hole.
18. Perform a firm but gentle double “simple knot,” performing a blind knot so as not to cut the tissue and having special care of not ligate neither interfering with the blood flow. **Note**: When necessary, hydrate the intestines with drops of sterile saline solution.
19. A second ligation is then performed at ∼1.5cm of distance from the first ligation using the same strategy described above (Fig 3B). However, in this case, perform a simple knot without closing.
20. At 0.5–1cm out of the second ligation, insert the syringe needle of a tuberculin syringe with 5×10^8^ *C. difficile* spores in 100µL of saline in the direction of the ligation and cross the ligation by inside the intestine the and close the knot with the syringe needle inside. *C. difficile* spores strain R20291 (CM210) were purified as was described previously [22].
21. Release the *C. difficile* spores inside the loop, keeping the pressure in the knot. This is performed to avoid inoculum loss and splashing that occurs when the ligated loop is injected directly.
22. Remove the syringe and close the ligation with a simple double knot. For steps 15–22, see S3 Video.
23. To perform the colonic loop, identify the regions to be ligated (Fig 3C) and repeat the points 15–22 in the proximal colon.
24. Carefully using anatomical forceps return intestines to the abdominal cavity. For steps 23 and 24, see S3 Video.
25. Close the peritoneum of the abdominal wall by a continuous or interrupted suture using silk suture.
26. Close the skin of the abdominal wall by continuous or interrupted suture using silk suture.
27. Remove mice from the isoflurane mask and from the heating pad and allow the mouse to recover from the anesthesia under a heat lamp. The awareness recovery usually takes 1– 2min. For steps 25–27, see S4 Video.
28. Apply postoperative analgesia when required according to the animal care protocol at your institution. Usually, the complete procedure for one mouse and one loop takes ∼15 min: and with 2 loops ∼10min.

**Fig 2.**
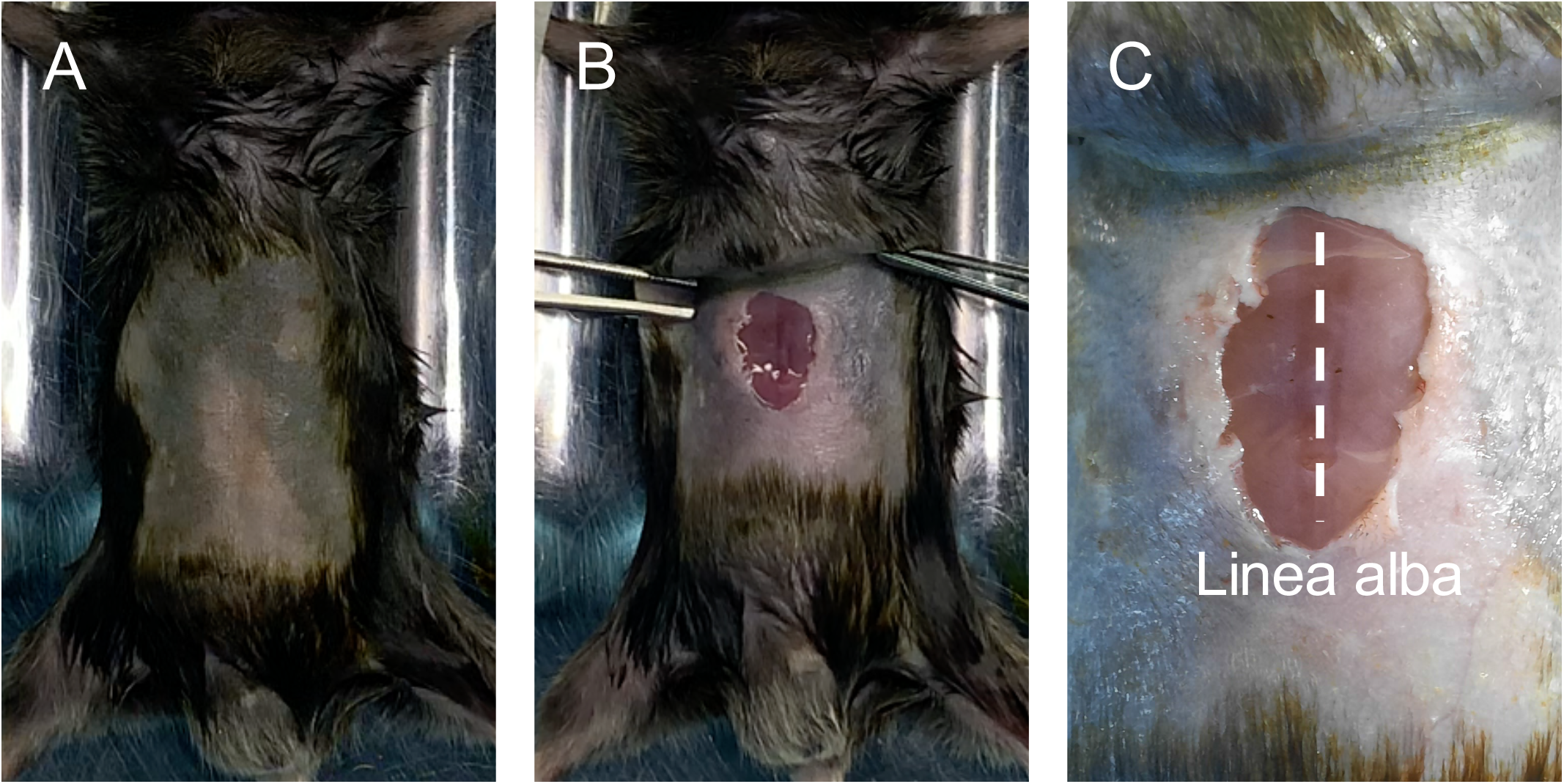
Schematic identification of linea alba. (A) The abdominal skin of the anesthetized mouse was disinfected with 70% ethanol, then was shaved, and the skin was cleaned with povidone-iodine. (B) The incision in the skin was performed parallel to the linea alba. (C) Identification of the linea alba as a semitransparent white line in the peritoneum.

**Fig 3.**
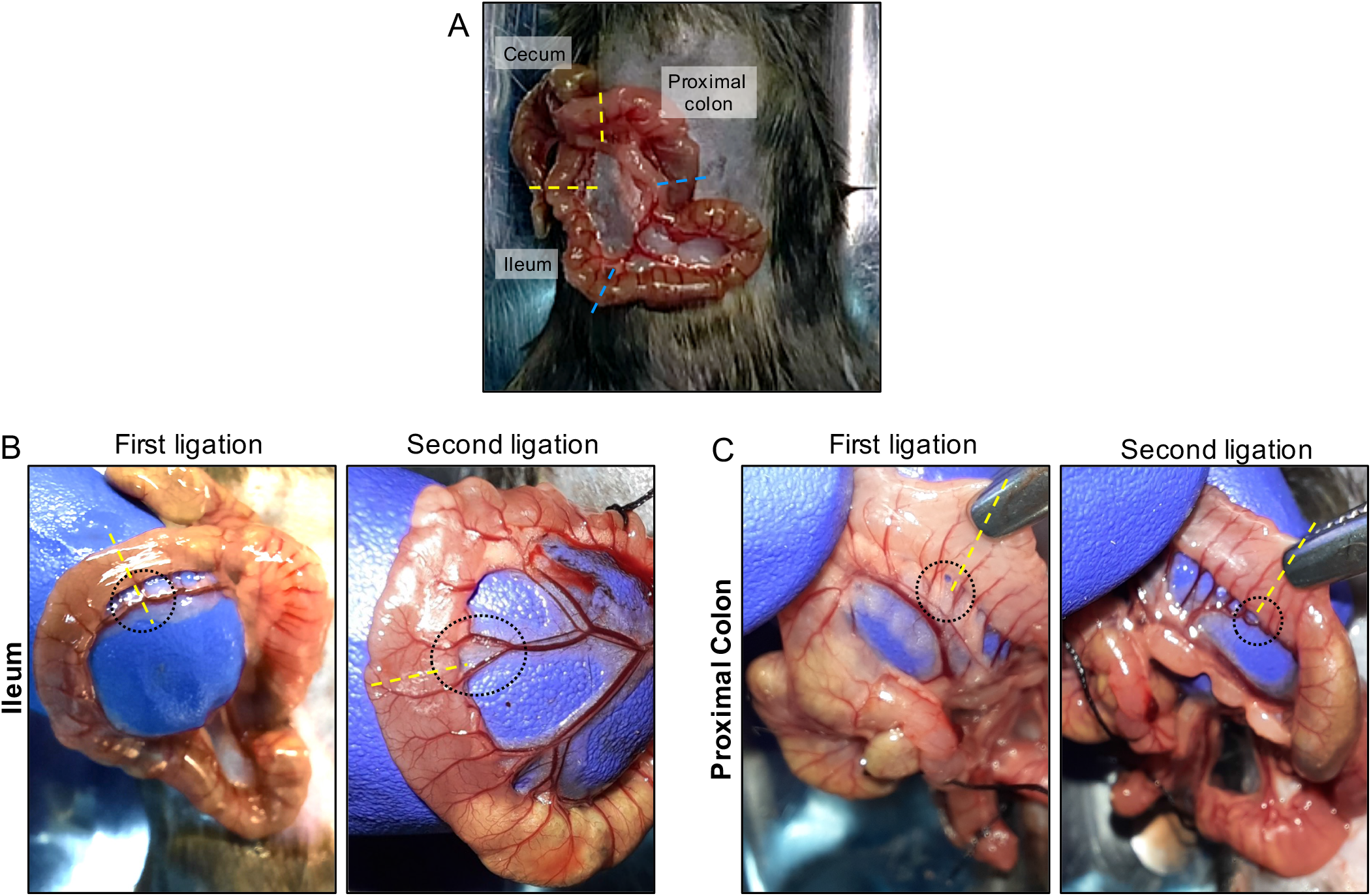
Identification of regions of interest to perform ligations between blood vessels. (A) Identification of ileum and colon using as reference the cecum. The regions of interest to be ligated are indicated with dotted lines. The yellow line and blue line denote the first and second ligation, respectively. Ligations are spaced ∼1.5 cm. The ligatures with surgical silk sutures were performed between the intestine and the blood vessels. The identification of regions of interest are shown in (B) ileum and (C) proximal colon. As a reference, the first ligation was performed close to the cecum.

Animals were kept in the cage for 5h with free access to water and close to a heat lamp. Animals were monitored every 30 min.

### Necropsy and tissue collecting

29. Depth anesthesia is induced by isoflurane inhalation, as is indicated above in step 2.
30. Check anesthetic depth by non-response to hind limb toe pinch.
31. Perform cervical dislocation by separating the vertebrae in the cervical area with a firm pinch to the neck using a rigid metallic tool and firmly pull the mouse from the tail. The separation of the skull and brain from the spinal cord is caused by anterior pressure applied in the skull base.
32. Using forceps, gently grab the skin and retract it up, so the abdominal organs and the ligated loops are not adjacent to the muscles.
33. Using scissors, open the abdominal cavity, cutting the skin and peritoneum. Extend the incision to visualize the intestines and the loops.
34. Remove the ligated ileum and colonic loop by cutting at ∼0.5mm from the outside of ligatures and put the intestinal loops in a petri dish.

For steps 33–34, see S5 Video.

### Fixing the tissues

35. In a biosafety cabinet, put drops of 1mL of PBS over a petri dish.
36. To fix the tissues, prepare a “fixation chamber”: we used a petri dish, but you can use any other tupperware with lid that you have available. Put inside a filter paper (extra thick blot paper). If the filter paper is larger than the used container, cut the filter paper with scissors o fit it inside the container.
37. Imbibe filter paper with the solution of 30% sucrose in PBS–4% paraformaldehyde enough to wet the entire filter paper without adding an excess of solution. Remove the excess of fixing solution from one edge using a 100–1000µL micropipette.
38. To remove the ligatures, cut the ligated loop as close as possible from the ligation. A liquid with gelatinous consistency comes out of it. Be careful with the handling because that liquid has a high concentration of *C. difficile* spores.
39. Put the scissor tip inside the lumen of the intestine and perform a longitudinal cut in the tissue to extend it.
40. Grasp the tissue from one end with the forceps and wash the tissue by immersion in the PBS drops for ∼20 immersions and repeat in 2–3 different drops of fresh PBS as is necessary for each tissue.
41. With anatomical forceps, grasp the opened intestinal tissues from a corner with the muscular layer downwards and the luminal side upwards. This can be identified because when the longitudinally cut tissue is grasped with the forceps from one end, it tends to recover its uncut shape, where the luminal side is inwards and the muscular side is outside. Using 2 pairs of forceps, stretch the tissues on the filter paper with the fixing solution.
42. Using a 100–1000µL micropipette, add fixing solution directly over the tissues. Repeat each ∼5min.
43. Let samples fixing for at least 15 min.
44. Using forceps, transfers each tissue to one independent 1.5mL microcentrifuge tube containing 1mL of 30% sucrose in PBS–4% paraformaldehyde solution, having care that the tissue is completely submerged in the fixing solution and there are no air bubbles in the tissue. Is common that some tissues should be folded in half so that it remains immersed in the solution. If the intestine has adipose tissue attached, it will tend to float. Therefore, it is recommended to remove the adipose tissue using scissors and forceps. Incubate the intestines in fixing solution overnight at 4° C.

For steps 36–44, see S6 Video.

### Immunofluorescence

#### Day 2

45. Using forceps, transfer the tissues to a new 1.5mL microcentrifuge tube containing 1mL of PBS. Perform this step carefully to avoid paraformaldehyde splashing. Incubate for ∼5 min a RT.
46. Wash the tissues. Using a 100–1000µL micropipette, remove the PBS and discard it to an autoclavable glass bottle, from now on, waste bottle. Add 1.0mL of PBS to the edges of the tube. And repeat one more time.
47. Using forceps put the tissues over a clean and sterile open petri dish, and using surgical scissors, cut a section of the tissues of ∼5mm × 5mm.
48. Using forceps, transfer the tissues to a 0.5mL microcentrifuge tube containing 150µL permeabilizing solution; PBS–0.2% Triton X-100 and incubated for 2h at RT.
49. Using a 20–200µL micropipette, remove the permeabilizing solution as much as possible from the walls of the tube and discard it in the waste bottle. Add 200µL of PBS to wash the samples and incubate for ∼3 min at RT in an orbital shaker at 60 RPM. Repeat 2 more times.
50. In the same tubes, incubate the tissues with 150µL of blocking solution; PBS–3% BSA for 3h at RT in an orbital shaker at 60RPM.
51. Using a 20–200µL micropipette, remove the blocking solution and discard it in the waste bottle.
52. Add 70–90µL of 1:1,000 chicken primary antibody anti-*C. difficile* spore IgY batch 7246 (AvesLab, USA) and 1:150 phalloidin Alexa-Fluor 568 (A12380 Invitrogen, USA); in PBS–3% BSA overnight at 4° C. This antibody does not immunoreacted with epitopes of vegetative cells neither with murine microbiota [20, 23]. After adding the antibody solution, check that there are no bubbles in the tube and that the tissue is completely submerged.

#### Day 3

53. Wash the tissues. Using a 20–200 µL micropipette, remove the primary antibody solution as much as possible from the walls of the tube and discard it in the waste bottle. Add 200µL of PBS to wash the samples and incubate for ∼3 min at RT in an orbital shaker at 60RPM. Repeat 2 more times.
54. In the same tube, incubate the tissue with 70–90 µL of 1:350 secondary antibodies goat anti-chicken IgY Alexa Fluor-647 (ab150175, Abcam, USA) and 4.5µg/mL of Hoechst 33342 (ThermoFisher, USA) and incubate for 3h at RT in an orbital shaker at 60RPM.
55. Wash the tissues 3 times as was described in step 53.

### Sample mounting

At this point is difficult to identify the luminal side and the muscular side of the tissues at the naked eye. However, sample mounting is essential to identify the tissue orientation. For this:

56. Using forceps, place the samples in a clean glass slide.
57. First, using a light -upright or -inverted microscope with 20× or 40× magnification coupled to epifluorescence with a blue filter to visualize Hoechst 33342 staining, orientate the tissues to put the liminal side up as follow.
  a. In the case of the ileum, villi can be visualized, and in the case of the colon, crypts are easy to identify. In both cases, on the other side, the muscular layer is seen.
  b. **Note 1**: if you are using upright microscopy, when you see the crypts or villi, keep the orientation of the tissue to the glass slide. If you use an inverted microscope, when you see crypts or villi, flip the tissue to the other side and put them in the glass slides. (Fig 4A and B).
  c. **Note 2**: During this process, don’t let the tissues dry because it causes autofluorescence. If samples begin to dry, add PBS with a micropipette.
58. Clean the coverslips and slides with 70% ethanol and towel paper.
59. Using towel paper removes the excess PBS from the edges of the tissues.
60. In a new clean glass slide, using a 2–20µL micropipette, put a drop of 5µL of fluorescent mounting medium for each tissue to be mounted.
61. Using forceps put the tissues over the drops of the mounting medium of the clean slide.
62. Using a 2–20µL micropipette, put 15µL of fluorescent mounting medium over the tissues having care of not to damage the tissue with the tip of the micropipette.
63. Put Scotch transparent tape (3M, USA) on the upper and lower edges of the coverslip, leaving half of the Scotch transparent tape on the coverslips and the other half free.
64. Put a coverslip over the samples, not allowing air bubbles to remain in the tissue.
65. With your fingers, fold the piece of Scotch transparent tape under the slide firmly.
66. Seal the remaining edges with Scotch transparent tape.
67. Store the samples at 4° C overnight, and then the samples are ready to visualize under confocal microscopy.
  a. Note: After mounting, we usually observe the samples as soon as possible and with no more than 1 week because sometimes samples begin to dry, making impossible the confocal visualization. If you need to store the samples for a longer time, you can store them in a wet chamber: in a tupperware with lid, put a layer of towel paper on the bottom and wet it with water (not in excess), then put a layer of parafilm or plastic wrap, and over it, you can put the samples and then close the cage.

**Fig 4.**
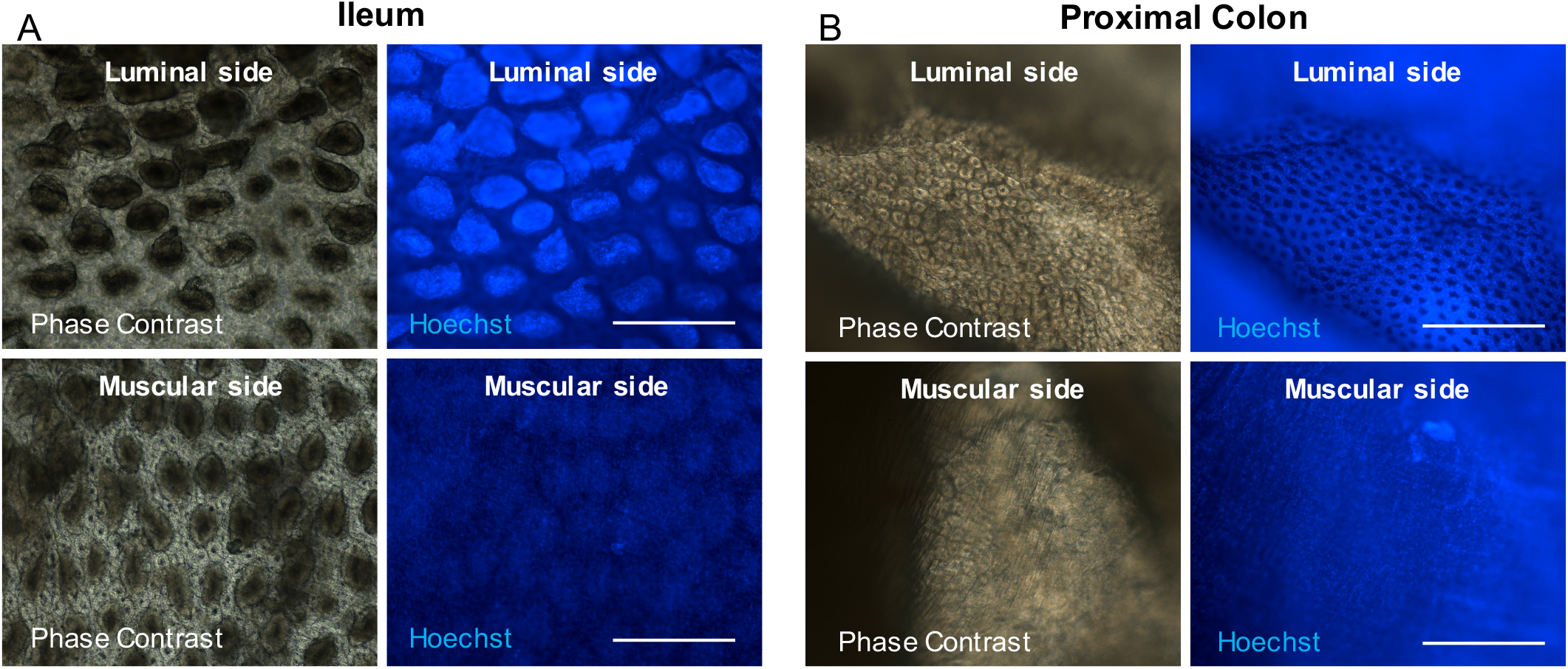
Tissue orientation under microscopy for mounting. Immunostained tissues were visualized under an upright light/epifluorescence microscopy to identify the tissue orientation to mount them with the luminal side up. Representative phase-contrast and Hoechst staining micrograph of (A) ileum and (B) proximal colon with the luminal or muscular side up. Scale bar 400µm.

For steps 57–67, see S7 Video.

### Confocal Microscopy

The confocal microscope Leica SP8 (Leica, Germany) of the Confocal Microscopy Core Facility of the Universidad Andrés Bello was used to acquire images. To evaluate spore adherence and internalization in the mice intestinal mucosa, images were acquired using the objective HPL APO CS2 40× oil, numerical aperture 1.30. For signals detection, three photomultipliers (PMT) spectral detectors were used; PMT1 (410–483) DAPI, PMT2 (505– 550), Alexa-Fluor 488, and PMT3 (587–726) Alexa-Fluor 555. Emitted fluorescence was split with dichroic mirrors DD488/552. Images of 1,024 × 1,024 pixels were acquired with 0.7-µm *z*-step size. Representative images were represented by three-dimensional (3D) reconstructions of intestinal epithelium using the plug-in 3D Projection of ImageJ software (NIH, USA). Villi and crypts were visualized by Hoechst and phalloidin signals.

### Quantification of spore adherence and internalization in the intestinal mucosa

Confocal images were analyzed using ImageJ. In the first place, we analyzed the spore adherence in the ileum mucosa in mice of 7-weeks-old, 1-, and 2-years-old. Representative images are shown in Fig 5A. Adhered spores were considered fluorescent spots in narrow contact with the actin cytoskeleton (visualized with F-actin). Adhered *C. difficile* spores were counted one-by-one using the plug-in Cell Counter or Point Tool of ImageJ. We observed that spore adherence varies between animals of each group and decreases according to increases the aging. The average spore adherence was ∼610, ∼571, and ∼427 spores, every 10^5^ µm^2^ in the ileum of mice with 7-weeks, 1-, and 2-years-old, respectively, with no significant differences between the groups (Fig 5B). We identified internalized spores using the plug-in Orthogonal View of ImageJ. Internalized spores were considered fluorescent spots inside the actin cytoskeleton in the three spatial planes (XY, XZ, YZ) [20, 24] (Fig 5A see magnifications XY and XZ). We observed that the spore internalization was on average of ∼0.5%, ∼0.3%, and ∼2.1% of the total spores in mice of 7 weeks old, 1- or 2 -year-old respectively, with a tendency to increase the spore entry in mice of 2-years-old compared to mice of 7-weeks-old (Fig 5C).

**Fig 5.**
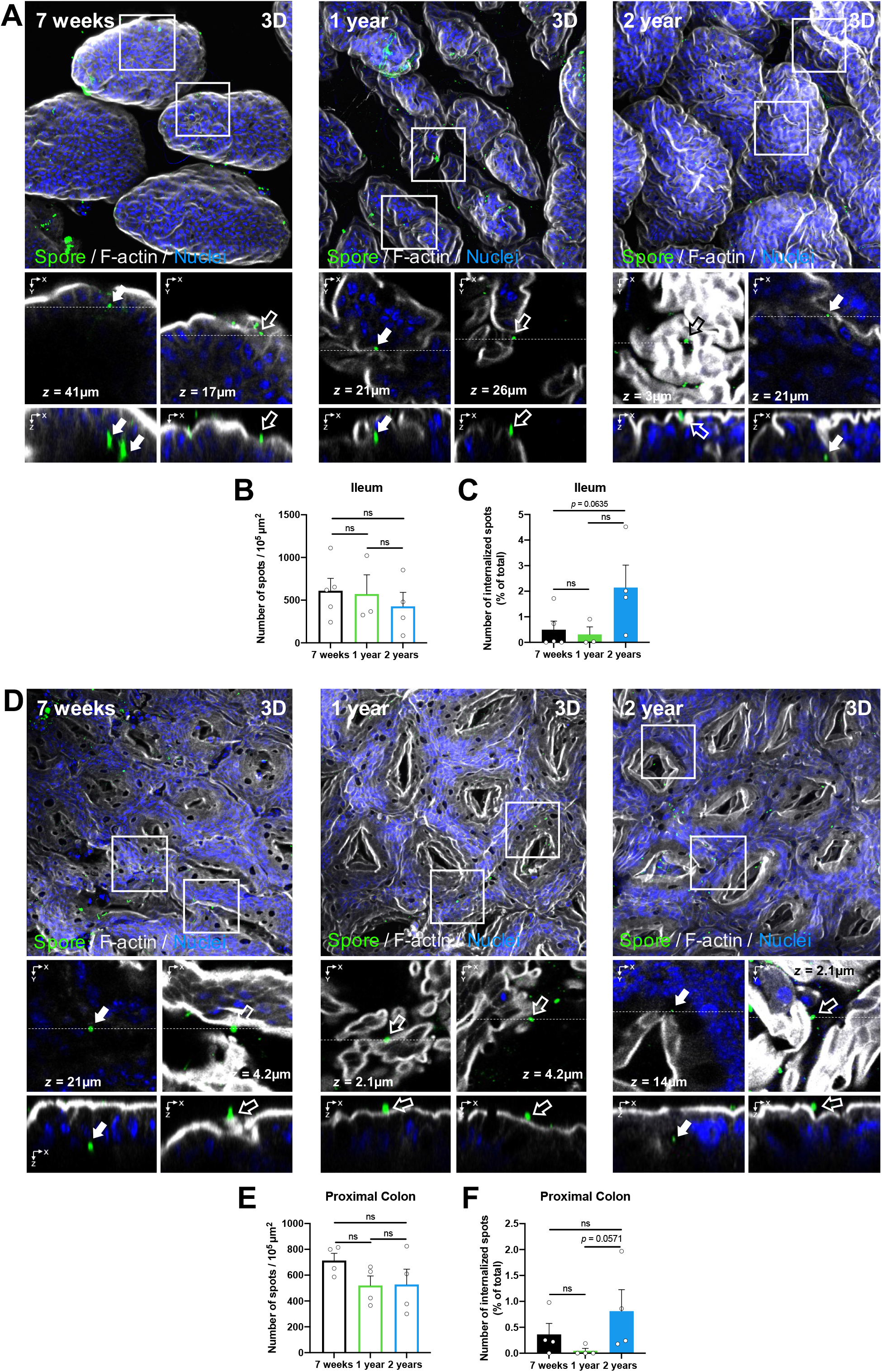
Visualization and quantification of adhered and internalized *C. difficile* spores in the ileum and colonic mucosa during aging. Representative confocal micrographs of (A) ileum and (D) colonic mucosa of the ligated loop. *C. difficile* spores are shown in green, F-actin in grey, and nuclei in blue (fluorophores colors were digitally reassigned for a better representation). The white arrow and empty arrow denote internalized and adhered *C. difficile* spores, respectively. Quantification of (B) adhered spots (spores) per 10^5^ μm^2^ and (C) percentage of internalized spots in the ileum or (E, F) colonic mucosa. Error bars indicate the average ± SEM. Scale bar, 20 μm. Statistical analysis was performed by two-tailed Mann–Whitney test; post-Dunn’s; test; ns, *p* > 0.05.

Using the same strategy, we analyzed spore adherence to the colonic mucosa. Representative images are shown in Fig 5D. We observed a decrease in spore adherence according to the aging increase. The spore adherence was on average of ∼713, 520, and 527 spores, every 10^5^ µm^2^ in the tissue of mice with 7-weeks-old, 1-year-old, and 2-years-old, respectively (Fig 5E). Then we observed that on average the 0.36, 0.05, and 0.81% of the total spores were internalized in the colonic mucosa, and we observed an increase in the spore internalization of mice of 2-years-old compared to mice of 1-year-old (*p* = 0.0571; Fig 5D magnifications XY and XZ and Fig 5F). Altogether, those data suggest that *C. difficile* spore adherence decreases according to increase aging, but the spore internalization increases in 2-years-old mice in both ileum and colonic mucosa.

## DISCUSSION

The intestinal ligated loop technique was described to our knowledge for the first time in 1953 in rabbits, [25], and since then has been widely used in several animal species such as rabbit [26-29], mouse, [30], rat [31], chicken [32] and pig [33] to study pathogenesis and the interaction with the host of bacteria such as *Clostridium perfringens* [34], *Vibrio cholerae* [35], *Listeria monocytogenes* [36], and *C. difficile* toxins TcdA and TcdB [26-29]. However, there is not a step-by-step protocol describing this technique coupled to confocal imaging to study the interaction of *C. difficile* with the intestinal mucosa.

In this work, we described a highly detailed protocol of a surgical procedure of intestinal ligated loop technique, including animal, anesthetize, opening the peritoneal cavities, perform the ligated intestinal loop, the inoculation with *C. difficile* spores, close the skin and peritoneum wall, monitoring the recovery of the mice, perform immunofluorescence of whole-mounted tissue against *C. difficile* spores, and staining of F-actin and nuclei, mounting of the sample, visualization of the sample by confocal microscopy, and finally the quantification of spore adherence and internalization in the ileum and colonic mucosa.

Using the intestinal ligated loop technique, here, we described that adherence of *C. difficile* spores to the ileum, and colonic mucosa is decreased in mice of 1-years-old and 2-years-old compared to 7-weeks-old mice. Also, that *C. difficile* spore entry is increased in 2-year-old mice. This finding, coupled with our recent observations that *C. difficile* spore entry is associated with R-CDI rates [20], may explain the increased R-CDI rates observed in elderly patients and in animal models [5, 6, 37].

During aging, several physiological changes occur in the intestinal mucosa that could affect spore adherence and internalization. In mice, it has been reported that a reduction of about 6-fold in the thickness of the colonic mucus layer in older mice compared to young mice [38], which enables a direct contact of bacteria with the intestinal epithelium and an increased bacteria penetration [38, 39]. Additionally, in human biopsies of older adults have been observed an increased intestinal permeability by a reduced transepithelial electric resistance compared to young humans [40] being those changes in the permeability to ions and not for macromolecules [41]. Recently we demonstrated that *C. difficile* spores gain access into the intestinal epithelial cells via pathways dependent on fibronectin-α_5_β_1_ and vitronectin-α_v_β_1_ [20]. Although fibronectin, vitronectin, and integrins α_5,_ α_v,_ and β_1_ are mainly located in the basolateral membrane [42, 43] we have shown that fibronectin and vitronectin are luminally accessible into the colonic mucosa of healthy young mice [20]. To date, whether these molecules are increased and/or become luminally accessible in aged intestines due to the reduced intestinal permeability and whether this contributes to spore adherence and entry to the intestinal mucosa remains to be elucidated.

We also recently shown that nystatin reduces the *C. difficile* spore entry *in vitro* and in the ileum but not into the colonic mucosa [20]. Nystatin is a cholesterol-chelating agent that disrupts cholesterol lipid raft required entry of pathogens dependent on caveolin and integrin [44, 45], suggesting that caveolin may be involved in *C. difficile* spore internalization. Results of this work showing that *C. difficile* spore internalization is increased with aging. Consequently, with this, had been reported that senescent cells had increased levels of caveolin-1, associated with higher rates of bacterial infection. For example, *Salmonella typhimurium* entry into senescent host cells that over-expressing caveolin-1 is increased compared with non-senescent cells, and the bacterial entry, depends on the levels of caveolin-1 expression [46]. However, those unknown changes in the aged intestinal mucosa, coupled with a reduced mucus thickness, could explain the increased *C. difficile* spore entry observed in the intestinal mucosa of older mice.

## Acknowledgments

The authors acknowledge to Nicolás Montes-Bravo for the help and dedication taking photographs and recording the videos shown in this work.

## Founding

This work was funded by FONDECYT Regular 1191601, 1151025, and by Millennium Science Initiative Program - NCN17_093, and Startup funds from the Department of Biology at Texas A&M University to D.P-S. Additional funding support to P.C-C from ANID-PCHA/Doctorado Nacional/2016-21161395. We certify that funding sources had no implication in the study design, data collection, analysis, and interpretation of data.

## Supporting information

**S1 Video. Mouse preparation for surgery** (steps 1–10). This video shows how to anesthetize the mouse, apply ophthalmic solution, disinfect, and shave the abdomen.

**S2 Video. Midline laparotomy** (steps 11–14). This video shown how to open the abdomen skin, identify the linea alba and open the peritoneal cavity.

**S3 Video. Procedure to ligate loops** (steps 15–24). This video shows how to identify the ileum and the proximal colon, remove fecal material from the section to be ligated, identify the sites to be ligated. Also, shown how to perform the ligations without interruption of the blood vessels and injection of *C. difficile* spores on ileum and colon.

**S4 Video. Midline laparotomy closure with suture** (steps 25–27). This video shows how to suture the abdominal wall and the abdominal skin with silk suture by continuous suture technique to close the incision and let mice recover from the procedure.

**S5 Video. Extraction of the ligated loop** (steps 33–34). This video shows how to extract the ligated loop in a euthanized mouse.

**S6 Video. Washing and fixing of extracted tissues** (steps 36–44). This video shows how to open and wash the infected ligated loops and the procedure of fixing with 30% sucrose in PBS–4% paraformaldehyde.

**S7 Video. Mounting of immunostained tissues for confocal microscopy** (steps 57–67). This video shows how to orientate the tissues to put the luminal side up of the ileum and the colon, the mounting using mounting medium, and sealing it with Scotch transparent tape.

