## Supplementary figures and images for "A ligated intestinal loop mouse model protocol to study the interactions of *Clostridioides difficile* spores with the intestinal mucosa during aging"

### Supplemental video 1

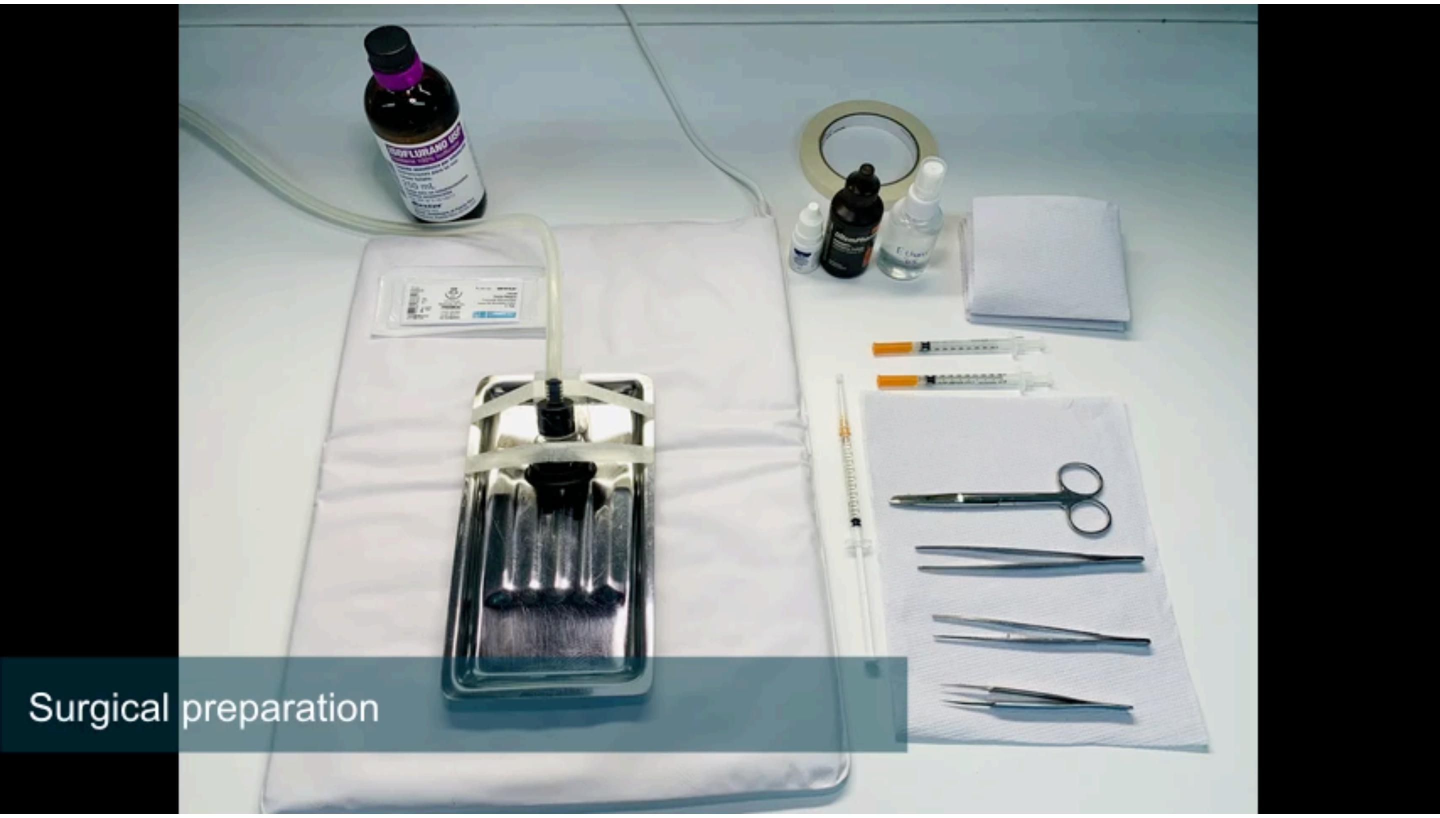

### Supplemental video 2

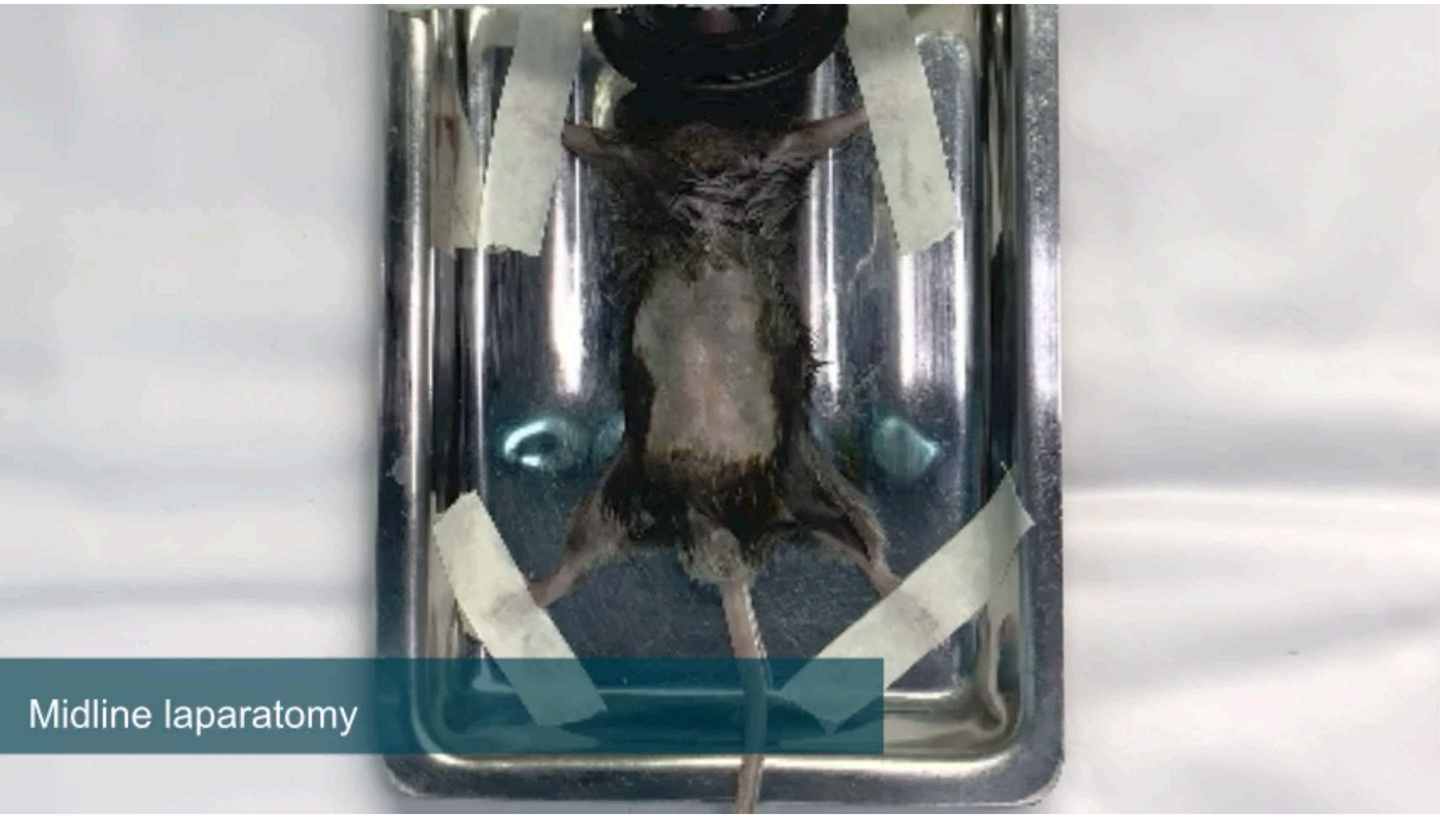

### Supplemental video 3

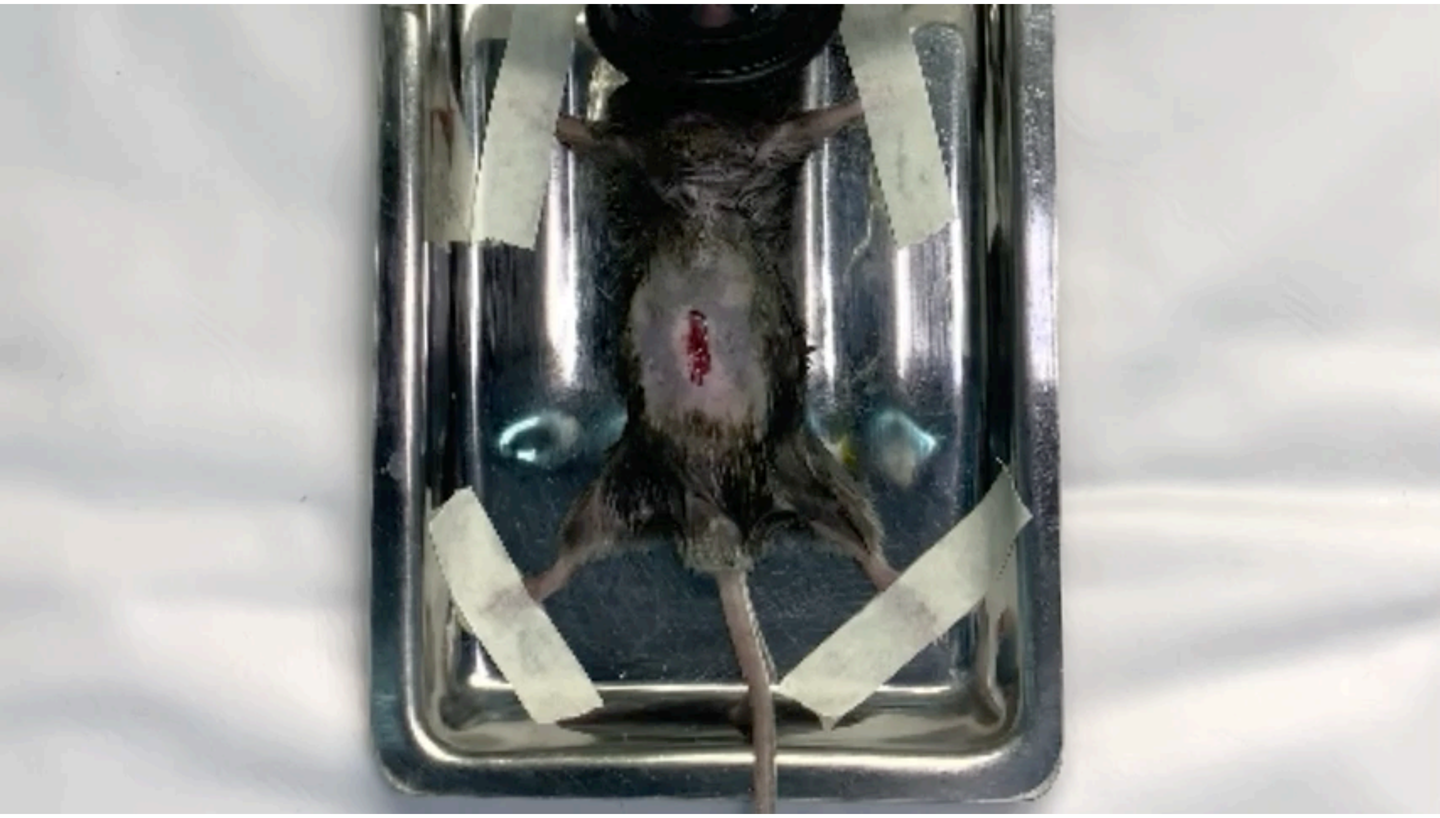

### Supplemental video 4

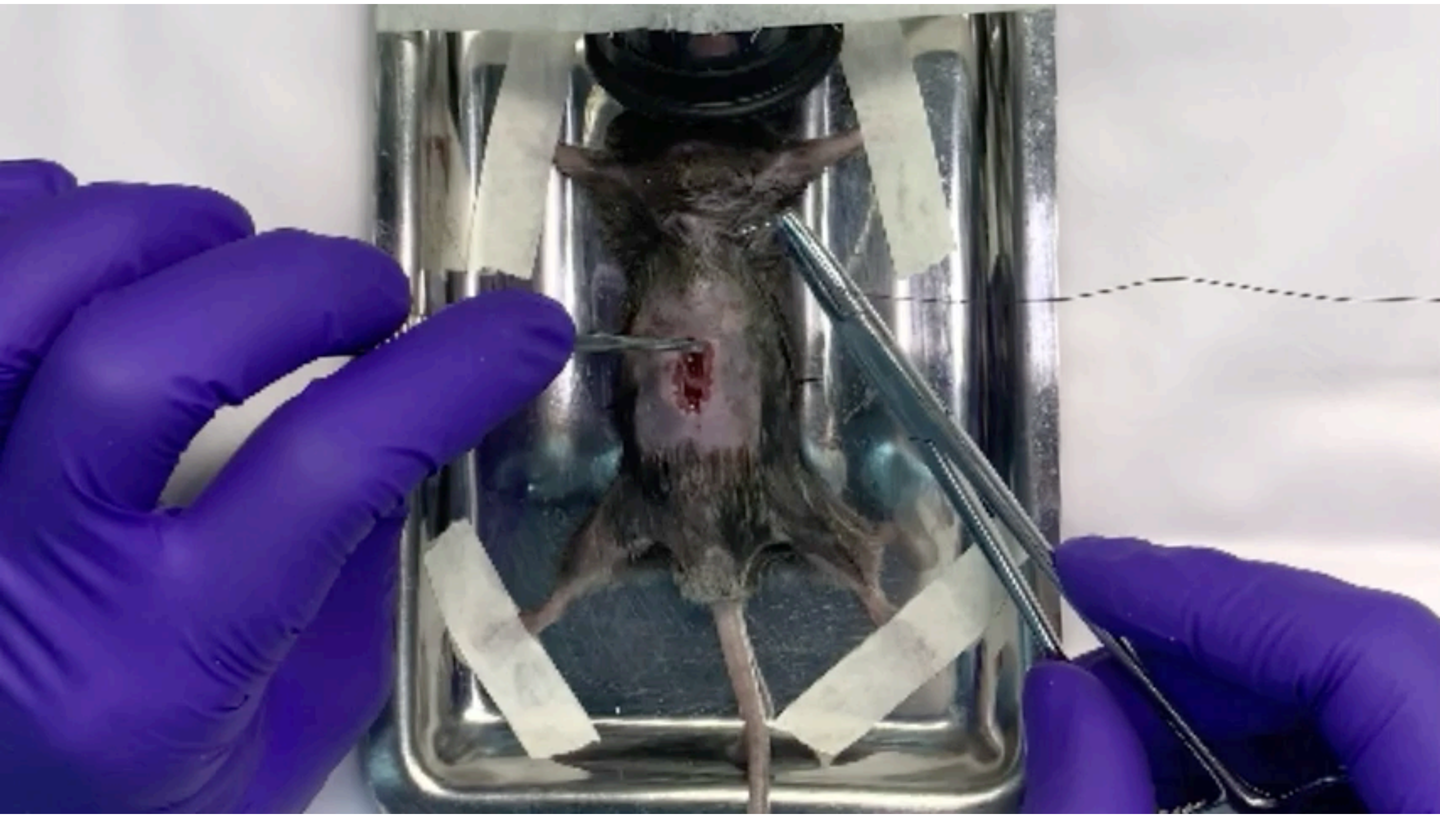

### Supplemental video 5

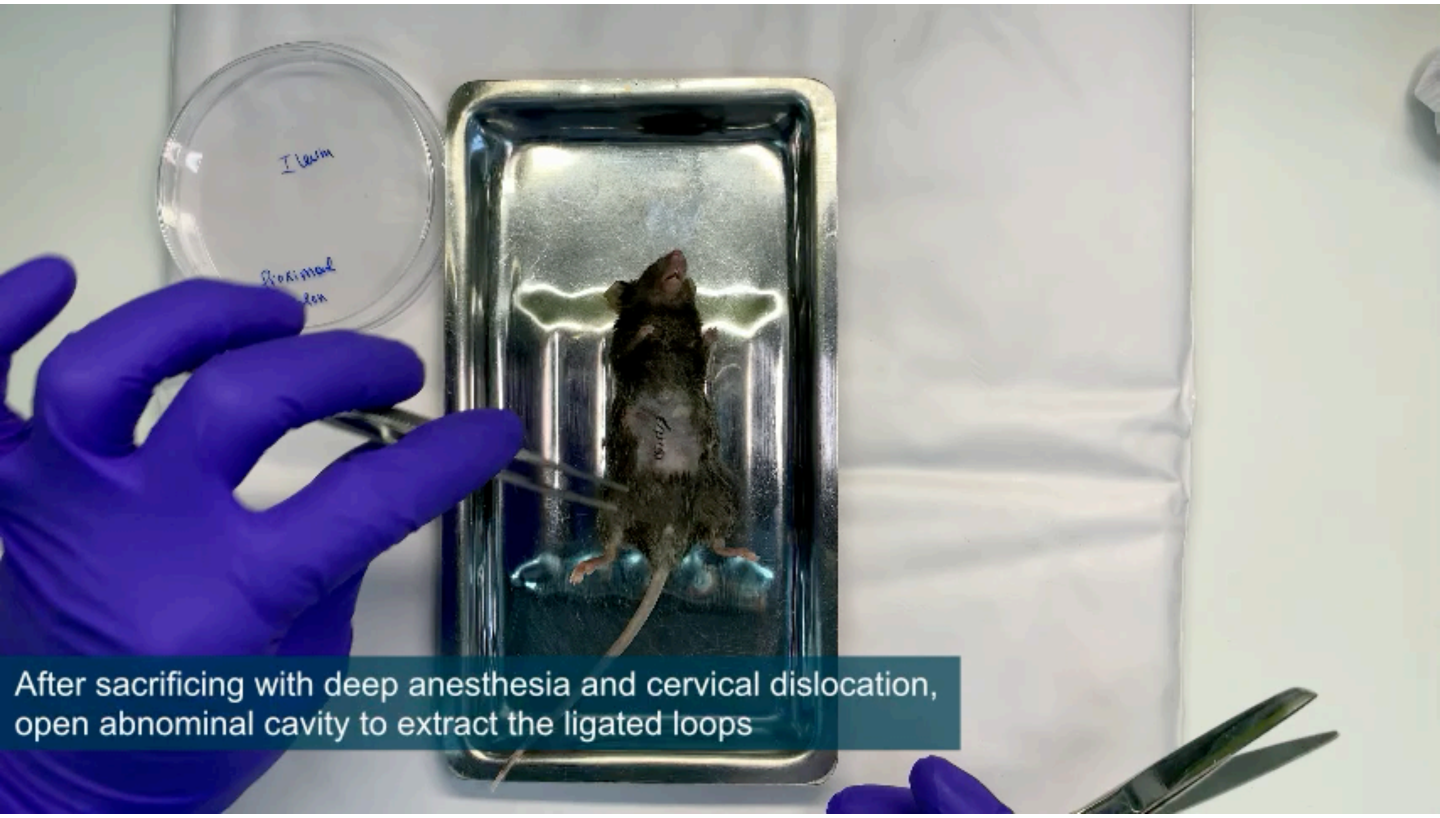

### Supplemental video 6

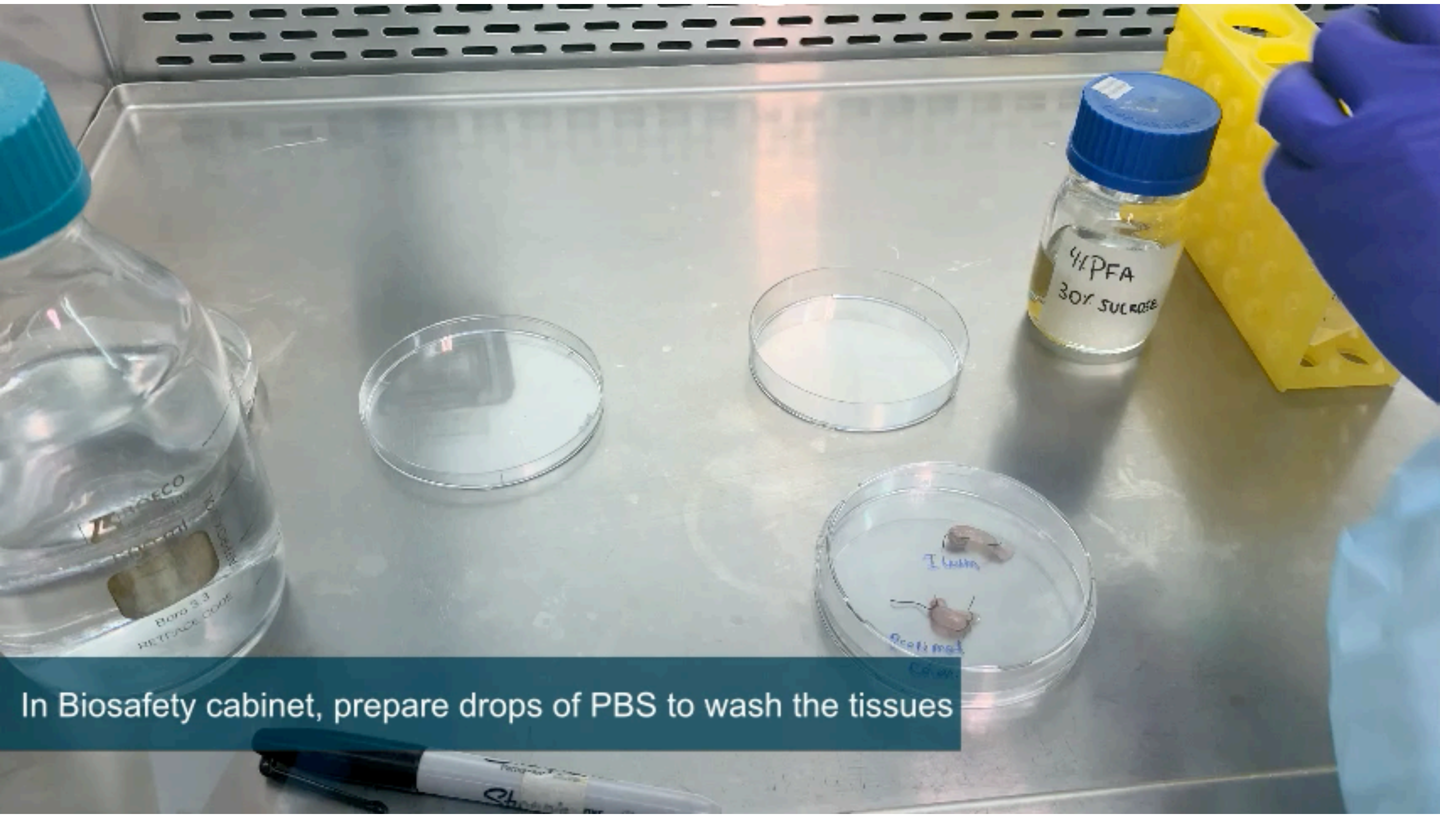

### Supplemental video 7

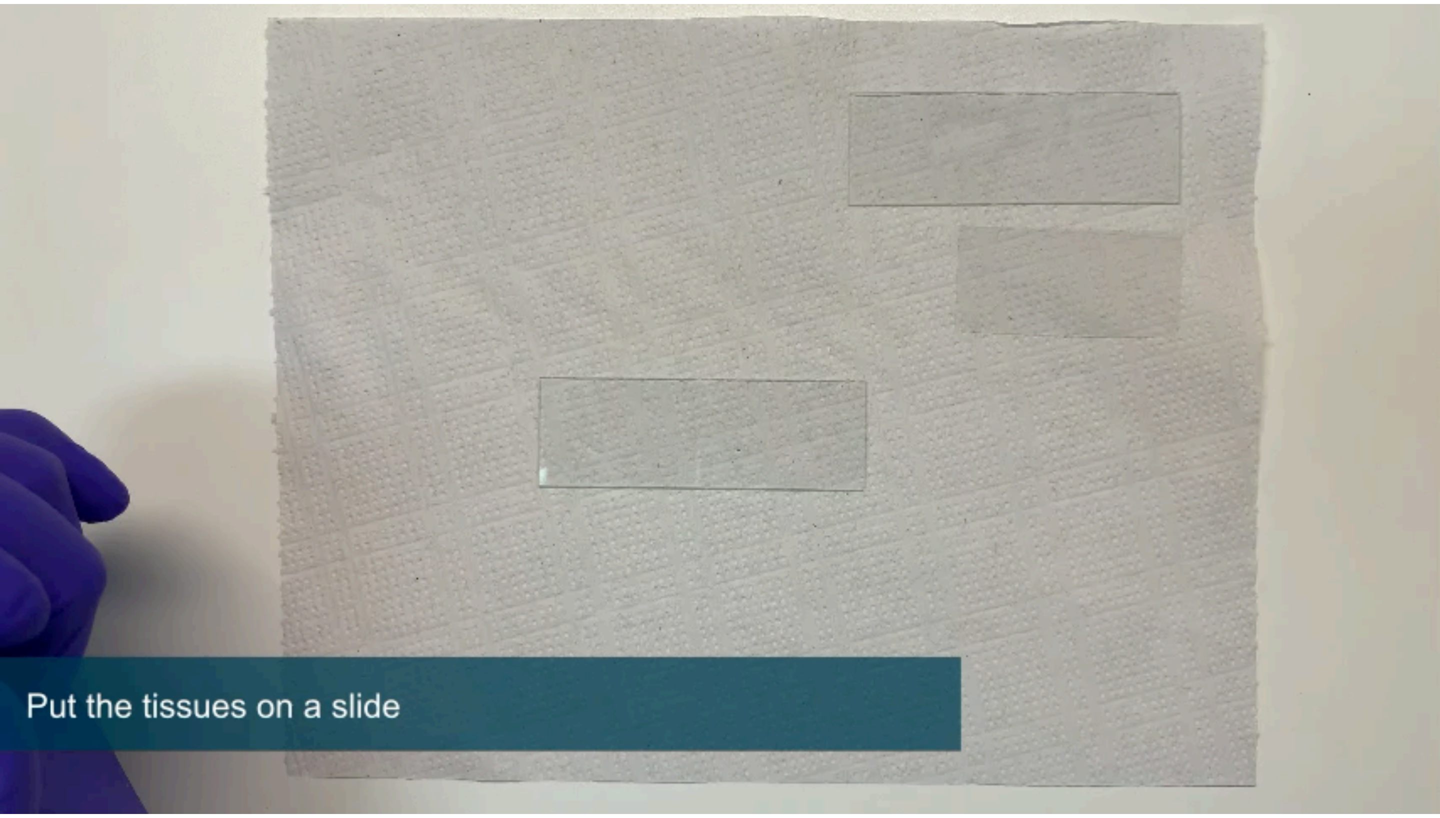
